# Oxidative Base Damage to Telomeres Sensitizes Cancer Cells to ATR Inhibition

**DOI:** 10.1101/2025.05.10.653274

**Authors:** Alex Garbouchian, Natalia Cestari Moreno, Aninda Dey, Patricia Opresko, Ryan Barnes

## Abstract

Targeted inhibition of DNA damage response proteins has received significant clinical attention owing to the success of PARP inhibitors. Due to the loss of the G1/S checkpoint, cancer cells are reliant on the G2/M checkpoint to cope with elevated DNA replication stress. We previously demonstrated a single induction of 8-oxo-guanine at telomeres in cancer cells was sufficient to induce replication stress, but was well tolerated at the cellular level. Here, we found inhibition of ATR, Chk1, or Wee1 after induction of telomere oxidative stress significantly induced genome instability and reduced cell viability. This occurred at doses markedly less than those required to increase instability in non-cancer cells. We determined the mechanism of this instability is due to cells progressing through S-phase with telomere damage and exiting G2-phase prematurely, prolonging their mitosis. This study demonstrates targeted oxidative base damage at telomeres can enhance the therapeutic efficacy of ATR inhibition in cancer.

## Introduction

Telomeres are nucleoprotein structures which prevent the DNA damage response machinery (DDR) from erroneously recognizing the ends of chromosomes as double-stranded DNA breaks (DSBs). Human telomeres consist of 5’ TTAGGG 3’ repeats on the order of several kilobases, which are bound by shelterin^1^. Shelterin is a 6-member protein group that binds both double-stranded (ds) and single-stranded (ss) telomeric DNA and organizes the chromosome end into a looped structure (t-loop) which obfuscates the free DNA ends^2^. While telomeres are critical for all cells, their relevance to proliferating cells is remarkably demonstrated in cancer, as every cancer cell engages a telomere maintenance mechanism. The most common mechanism is telomerase upregulation (TEL), but approximately 10-15% of cancer use a homologous recombination based, alternative lengthening of telomeres (ALT) pathway instead.

Seminal work has shown that the removal of shelterin member TRF2 from mammalian cells disrupts telomere function, leading to t-loop dissolution and chromosome fusions^3^. Removal of TRF1 or POT1 instead, increases DNA replication stress at telomeres specifically^4^. This is in part, thought to be due to the inherent difficulty in replicating the repetitive telomere sequence, which requires the recruitment of DNA replication and repair factors like PARP1 and BLM, mediated by TRF1^5, 6^.

Replication stress is broadly defined as the slowing or stalling of DNA replication forks. Approximately 2 decades ago, work from the Bartek and Halazonetis groups postulated that replication stress in precancerous cells was a driver of tumorigenesis due to increased genomic instability^7, 8^. We now know that stressed replication forks can collapse due to exhaustion of replication factors, nuclease digestion, and replisome disassembly^9^. Thus, cells require increased time following replication stress to complete DNA synthesis so that they do not enter mitosis with under-replicated DNA, which causes chromosomal instability when cells divide.

ATR is the principal regulator of the replication stress response (RSR) in mammalian cells^10^. Stalled replication forks generate considerable ssDNA which is protected by RPA binding and the accumulation of this RPA coated ssDNA is the signal for ATRIP, TOPBP1, and others to activate ATR. Once activated, ATR phosphorylates a variety of targets including its main effector kinase, Chk1, to inhibit cell cycle progression from S and G2 to mitosis, suppress origin firing, and recruit DNA repair and replication factors^10, 11^. Unlike the G1/S checkpoint, the S and G2/M checkpoints are rarely mutated in cancer, as cancer cells rely on them to mitigate the elevated genomic instability and replication stress they experience^12^. Consequently, inhibitors of ATR and Chk1 have received increased clinical attention in recent years, with at least five drugs in Phase II trials, and one in Phase III^13^. These include VE822 and AZD6738, the first two ATRi to enter clinical trials, which are very specific ATP competitive inhibitors of ATR kinase activity, and are used in the clinic in combination with DNA damaging agents.

In addition to replication stress, cancer cells display elevated levels of reactive oxygen species (ROS), leading to oxidative stress^14^. While beneficial at slight elevations, cancer cells must maintain a balance between ROS and their antioxidant defenses to maintain viability^15^. Exploiting this tightrope cancer cells need to walk is another therapeutic option being pursued^16, 17^. Our group has previously shown that chronic oxidative stress at telomeres in the form of singlet oxygen mediated 8-oxo-guanine (8oxoG) formation significantly reduces cancer cell growth, and this coincided with increased replication stress and genome instability^18, 19^. However, cancer cells were largely unaffected by a single induction of telomeric 8oxoG, but did display slight increases in replication stress markers. Non-diseased cells senesced after a single treatment, and this senescence was dependent on DNA replication following 8oxoG induction^20^. Together our findings showed telomeric 8oxoG induces replication stress in dividing cells, but cancer cells were largely refractory to a single burst of this damage alone.

In this study, we sought to determine if oxidative base damage to telomeres would sensitize cancer cells to low-dose ATRi. Remarkably, ATR inhibition with VE822 or AZD6738 induced genome instability following induction of 8oxoG at telomeres in multiple cancer cell lines, coinciding with reduced cell viability. Non-cancer cells with intact p53 signaling were additionally sensitized to telomere replication stress by ATRi, but at doses 2-10 fold higher than cancer cells. We found ATRi and telomere 8oxoG induced genome instability was dramatically elevated when cells were damaged at the G1/S-phase boundary or mid-S-phase, confirming replication stress is responsible for this phenotype. Permitting cells increased time in G2 during these treatments however, rescued genome stability. Together, our data highlight that while telomeres are a very small portion of the genome, when they experience replication stress, this must be properly managed by the RSR or cells prematurely exit G2-phase and produce mitotic errors, leading to reduced viability.

## Results

### Telomeric 8oxoG Induced Replication Stress Requires ATR and Chk1 to Maintain Genome Stability

Our prior work demonstrated DNA synthesis through 8oxoG in telomeres disrupts DNA replication and causes telomere replication stress^18-20^. This led us to postulate that restraining the replication stress response (RSR) through ATR or Chk1 inhibition, would sensitize cancer cells to telomere replication stress, as cancer cells are more reliant on ATR signaling for survival (Figure S1A). Consistent with our previous studies, U2OS (ALT) and HeLa LT (TEL) FAP-TRF1 cells treated with MG2I photosensitizer dye and 660 nm light to produce 8oxoG at telomeres had no increase in genome instability as evaluated by counting micronuclei (MN) (Figure 1 and Figure S1B). However, when ATR inhibitors VE822 or AZD6738 were added after dye and light treatment, we observed significant and dose-dependent increases in MN for both cell lines (Figure 1A-D). Inhibiting the main target of ATR, Chk1, using a specific inhibitor (LY2603618) after 8oxoG induction also generated significant genome instability in both cell lines (Figure 1E, F). Importantly, for both cell lines and all three drugs, lower ATRi/Chk1i doses induced MN only after dye and light treatment, without increasing MN on their own. This demonstrates telomeric 8oxoG induces considerable replication stress, that while normally tolerated in cancer cells, elevates genome instability when the RSR is inhibited.

**Figure 1:**
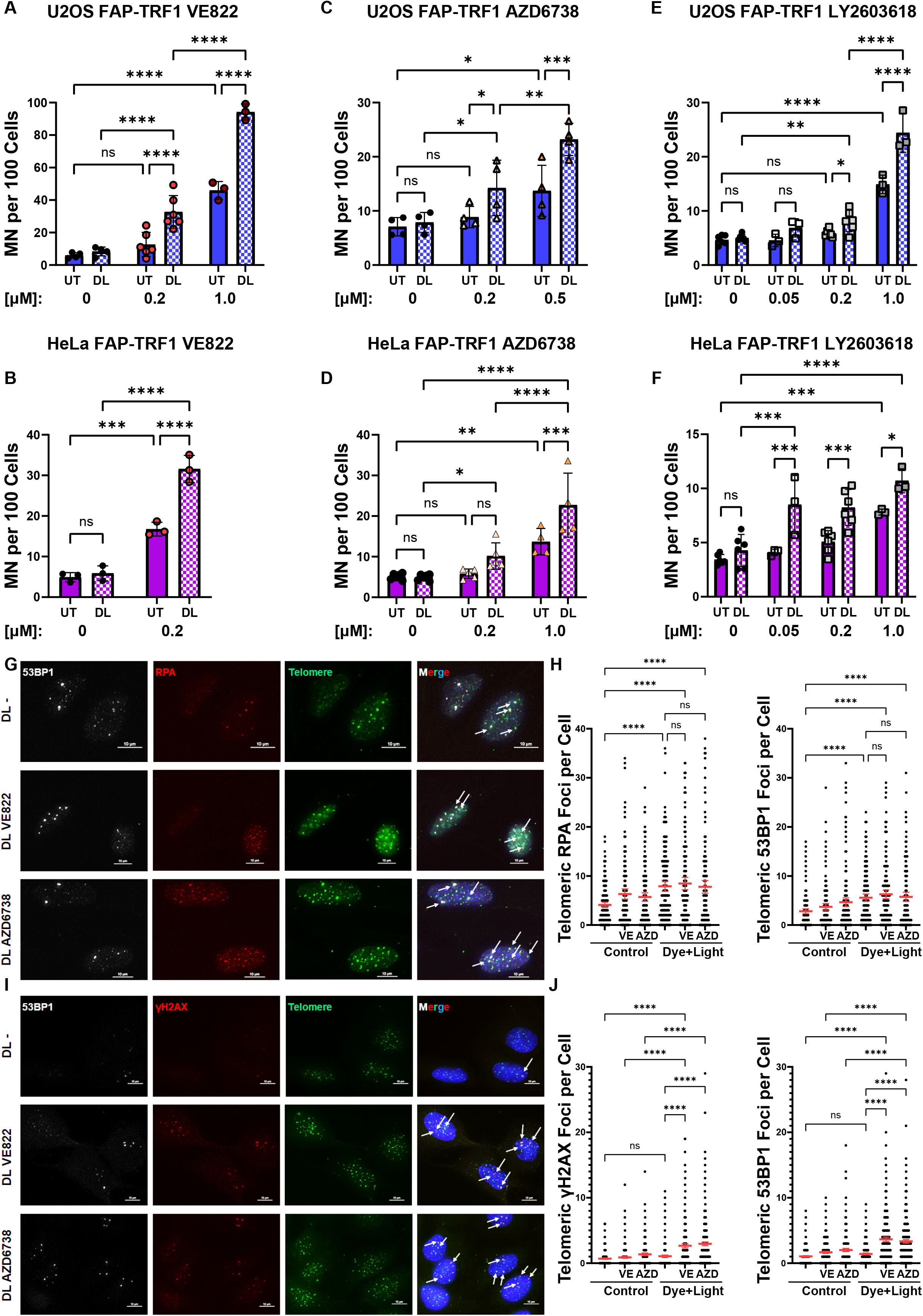
Telomeric 8oxoG Induces Genome Instability in RSR Inhibited Cancer Cells. (A-F) Micronuclei (MN) were quantified in U2OS and HeLa FAP-TRF1 cells 24 hours after 5 minute dye and light treatment. Where indicated, ATRi VE822 (A, B), ATRi AZD6738 (C, D), or Chk1i LY2603618 (E, F) were added after 8oxoG induction. Data are mean ± SD. Each point is a biological replicate. (G-J) U2OS FAP-TRF1 cells were treated with dye and light for 5 minutes, and recovered for 3 (G, H) or 24 hours (I, J). Where indicated, 0.05 μM VE822 or 0.2 μM AZD6738 were added after 8oxoG induction. Numbers below graphs are drug concentration in μM. Data are the mean and 95% confidence interval. Points are individual cells (150-400) from 3 biological replicates. All data analyzed by 2-way ANOVA.

Indeed, we found telomeric 8oxoG induction significantly increased RPA foci at telomeres 3 hours after treatment. Increased levels of RPA on chromatin are the principal signal for ATR activation, demonstrating 8oxoG is sufficient to disrupt telomere replication (Figure 1G, H). However, addition of ATRi did not further increase RPA levels, suggesting 8oxoG alone induces considerable replication stress. At this same timepoint, we find increased telomeric 53BP1 foci, indicating telomere dysfunction, but again ATRi did not increase this over 8oxoG induction. 24 hours after treatment, cells resolve the telomere DDR (53BP1 and γH2AX foci) caused by 8oxoG (Figure 1I, J). However, with ATRi this telomere DDR persists demonstrating telomeric 8oxoG requires ATR signaling to restore telomere stability and function.

We confirmed the genome instability phenotype in two other cancer cell lines (SAOS (ALT) and LOX (TEL)), and found both display increased MN when ATR is inhibited following 8oxoG induction at telomeres (Figure S1E, F). This indicates the induced genomic instability is not specific for cells which use telomerase or ALT for telomere maintenance. We also confirmed 0.2 μM VE822 and AZD6738 was a sufficient dose to suppress ATR signaling even in response to UVC (Figure S1 C). Since ATM is closely related to ATR, we tested if its inhibition following telomeric 8oxoG induction affected genome stability. However, even at a dose which inhibits hydrogen peroxide induced signaling, ATMi did not alter MN in U2OS or HeLa cells (Figure S1D, G, H). Finally, we tested a second Chk1 inhibitor (AZD7762), however this drug also inhibits Chk2^21^. We observed increases in MN in both HeLa and U2OS cells after dye and light treatment, but the effects were modest compared to LY2603618 which is highly specific for Chk1 (Figure S1I, J). In total, we found the replication stress generated by 8oxoG enriched telomeres is sufficient to require the RSR for genome integrity, sensitizing cancer cells to low ATR/Chk1 inhibition.

### ATR Maintains Cell Viability Following Telomere Oxidative Base Damage

In our previous studies we found chronic formation of 8oxoG at telomeres reduced cancer cell viability, but an acute treatment was viable even with increasing light exposure times (more 8oxoG)^18, 19^. Following our genome instability results, we wondered if ATRi after telomere 8oxoG induction would reduce cancer cell viability. For these experiments, we treated U2OS and HeLa cells with dye and light, and low dose VE822 and AZD6738 which alone is not cytotoxic (Figure S1). Consistently, we found induction of 8oxoG at telomeres sensitized cancer cells to low dose ATRi as measured by counting cells 4 days after treatment (Figure 2A-D). We confirmed this result with clonogenic survival experiments, and again found reduced cell viability with the combination of ATRi and telomeric 8oxoG (Figure 2E, F). Surprisingly however, when we performed Annexin V and propidium iodine staining of these cells, we observed no increased in apoptosis or necrosis after 8oxoG induction and ATRi (Figure 2G, H and S2). This is similar to our observations chronically treating cancer cells with dye and light, in which we observed significant reductions in population doubling, but no obvious change in cell death

**Figure 2:**
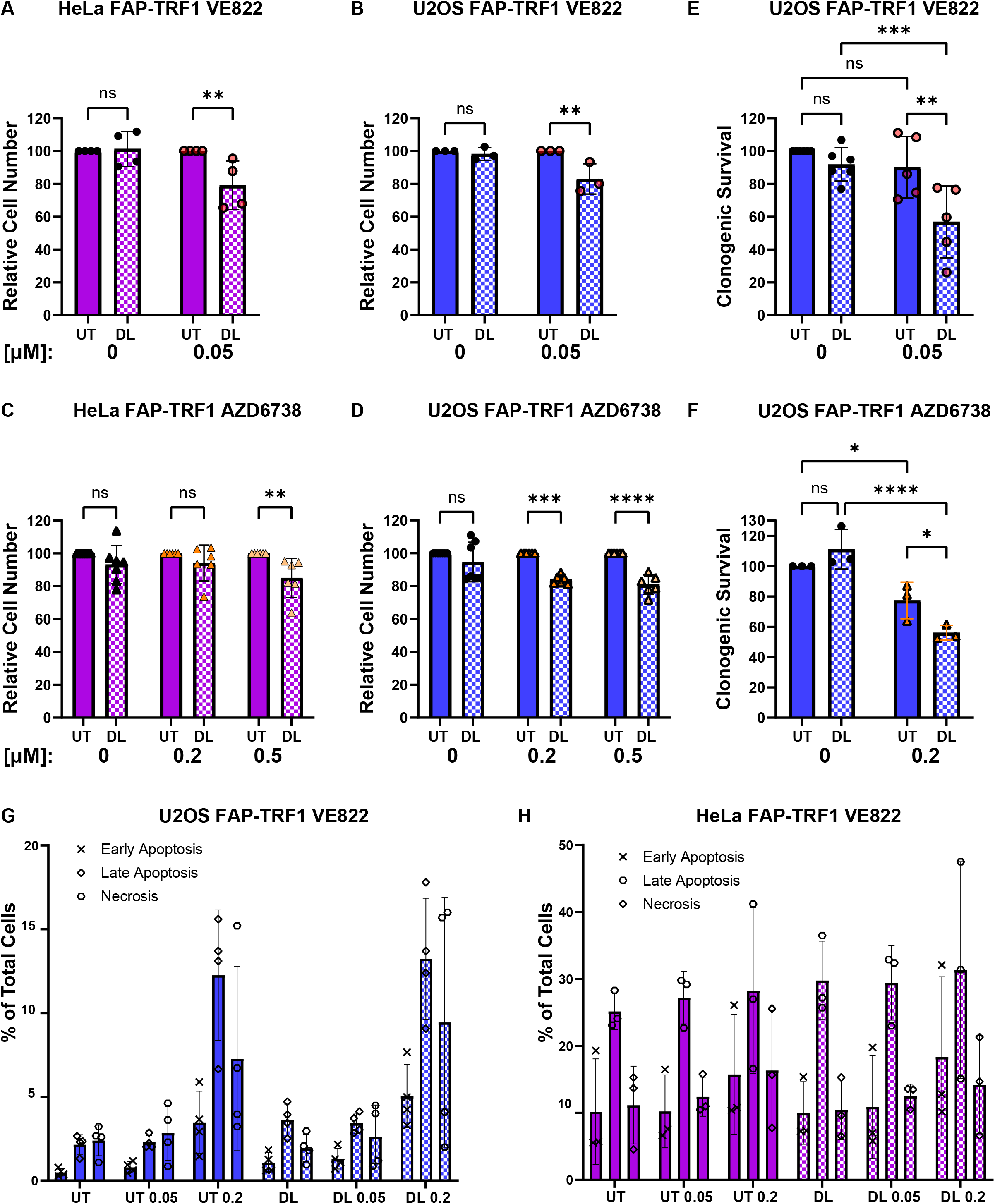
ATR Maintains Cell Viability Following Telomere Oxidative Base Damage. (A-D) HeLa and U2OS FAP-TRF1 cells were treated with dye and light for 5 minutes, and recovered for 4 days before counting. Where indicated VE822 (A,B) or AZD6738 (C,D) were added after 8oxoG induction. Data are normalized to the UT condition for each drug dose. (E, F) U2OS Fap-TRF1 cells were treated with dye and light for 5 minutes, and recovered for 7-10 days in the presence or absence of VE822 (E) or AZD6738 (F). Data are normalized to UT no drug. (G,H) Cells were treated and recovered as in A,B and then stained with Propidium Iodine and Annexin V to measure apoptosis and necrosis. Numbers below graphs are drug concentration in μM. All data are mean ± SD. Each point is a biological replicate. All data analyzed by 2-way ANOVA.

### Oxidative Telomere Damage Does Not Sensitize Non-Cancerous Cells to Low Dose ATRi

Our previous work showed telomeric 8oxoG induced senescence in non-diseased, otherwise normal, cell lines in a manner dependent both on p53/ATM and DNA replication following damage^20^. To understand how non-cancer cells would respond to ATRi after 8oxoG formation at telomeres, we compared RPE-1 FAP-TRF1 cells to U2OS and HeLa FAP-TRF1 cells treated with dye and light and 0.05 μM VE822 (Figure 3A, B). At this very low dose, both cancer lines displayed genome instability following 8oxoG induction, but not ATRi alone. RPE cells however, while displaying the small MN increase with dye and light treatment we described previously, showed no change with VE822. Consistently, RPE colony survival after telomeric 8oxoG induction was reduced as we previously showed, and this was unaffected by 0.05 and 0.2 μM VE822 (Figure 3C)^20^. At 0.5 μM VE822, a dose cytotoxic to U2OS and HeLa (Figure S1A), we observed a significant reduction in colony survival compared to the no drug control after 8oxoG induction.

**Figure 3:**
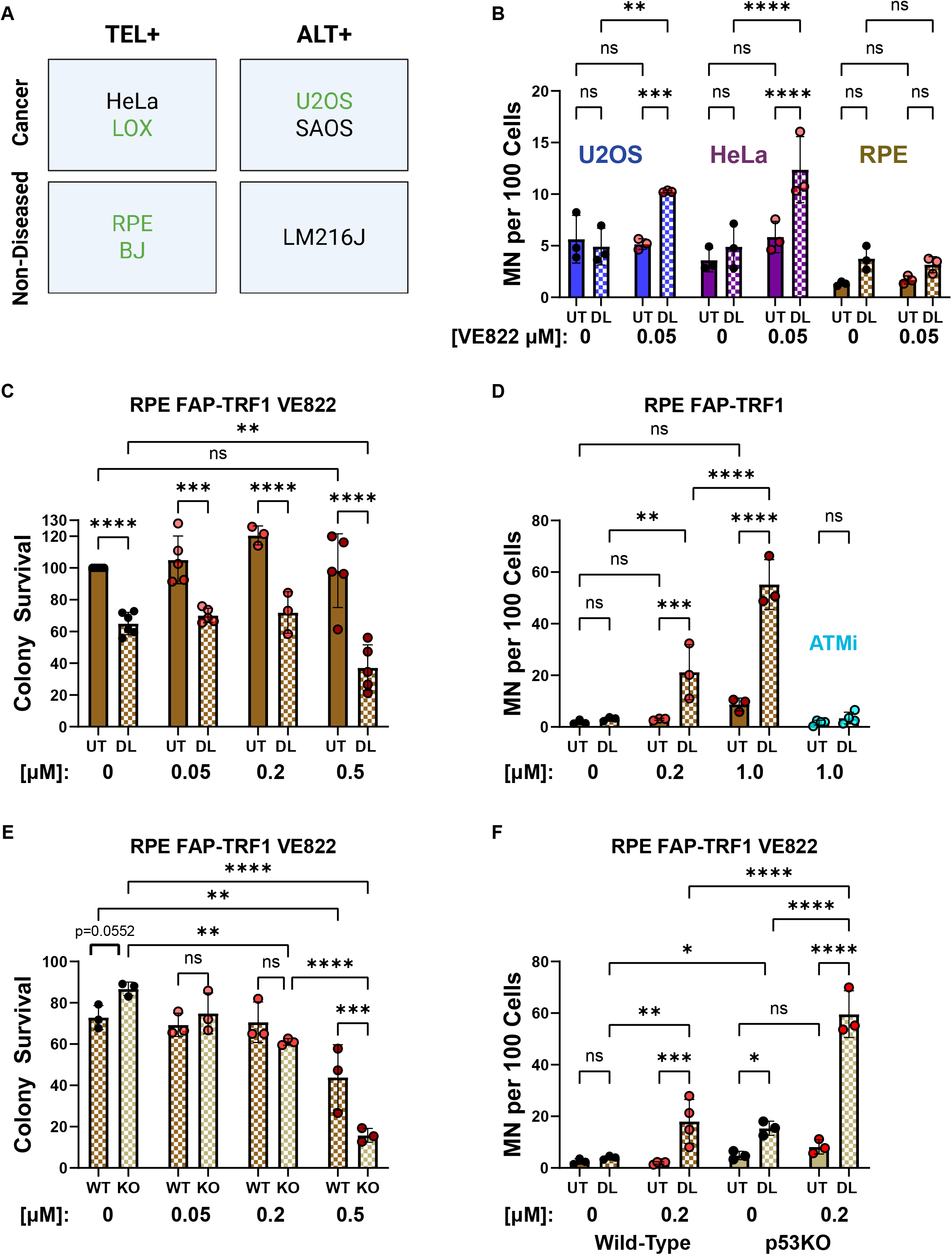
Oxidative Telomere Damage Does Not Sensitize Non-Cancerous Cells to Low Dose ATRi. (A) Table of cell lines used in this study, denoting those derived from cancer or non-diseased tissue, and the telomere maintenance mechanism. Green lines have wild-type p53 gene. (B) FAP-TRF1 cells were treated with dye and light for 5 minutes, and recovered 24 hours with or without 0.05 μM VE822. (C) RPE FAP-TRF1 cells were treated with dye and light for 5 minutes, and recovered as indicated with VE822 for 7-10 days. Data are normalized to UT no drug. (D) RPE FAP-TRF1 cells were treated with dye and light for 5 minutes and recovered with VE822 (red points) or ATMi for 24 hours. (E) Wild-type and p53ko RPE cells were treated with dye and light for 5 minutes, and recovered as indicated with VE822 for 7-10 days. Data shown are dye and light treated samples, normalized to UT no drug for each line. (F) Wild-type and p53ko RPE cells were treated with dye and light for 5 minutes, and recovered with or without 0.2 μM VE822 for 24 hours. Numbers below graphs are drug concentration in μM. All data are mean ± SD. Each point is a biological replicate. All data analyzed by 2-way ANOVA.

Since higher doses of VE822 further sensitized RPE cells to telomere replication stress, we treated them with 0.2 and 1.0 μM VE822 and quantified MN (Figure 3D). Despite no additional change in colony survival, 0.2 μM VE822 treated cells had a significant increase in MN after 8oxoG induction, however the total number of MN was still much lower than cancer cells. 1.0 μM VE822 dramatically raised MN after dye and light treatment, but also the background level. With 0.2 μM AZD6738 RPE cells showed no increase in MN or change in colony survival, but higher doses elevated MN and reduced colony survival (Figure S3A, B). As with U2OS and HeLa, ATMi did not alter MN after dye and light treatment and ATRi after 8oxoG induction did not induce apoptosis (Figure 3D and S3C).

While there are countless differences between non-diseased and cancer cells, we wondered if fully functional p53 signaling was allowing RPE cells to tolerate ATRi better than cancer cells. We repeated clonogenic survival experiments with wildtype and p53ko RPE cells using the same increasing doses of VE822. Remarkably, we found increasing ATRi reversed the phenotypic differences previously observed for wild-type and p53ko RPE cells. As we previously reported^20^, without ATRi, p53ko cells were more resistant to telomeric 8oxoG, but this was nullified with 0.05 μM VE822 (Figure 3E). 0.2 μM VE822 significantly reduced p53ko cell survival compared to no ATRi control, and at 0.5 μM, p53ko RPE had a significant reduction in survival compared to wildtype cells. Consistent with this, when p53ko RPE cells were treated with dye and light and ATRi, they displayed a dramatic increase MN, significantly greater than wildtype cells (Figure 3F). Similar results were observed for BJ FAP-TRF1 wildtype and p53ko cells, which are another non-diseased cell line pair (Figure S3D, E). We also treated LM216J cells with VE822 following 8oxoG induction, and found an increase in MN (Figure S3F). This line is not derived from cancer, but is SV40 E6/E7 transformed, abrogating both p53 and p16 signaling pathways, and uses ALT for telomere maintenance^22^. Collectively, our data suggest a conserved mechanism exists for preventing genome instability following telomere replication stress in cancer and non-cancer cells alike, and the tumor suppressor p53 has a critical role in this mechanism.

### DNA Replication is Hypersensitive to Telomeric 8oxoG

Telomeres represent a small (0.02%) portion of the genome, which likely means when asynchronous FAP-TRF1 cells are treated with dye and light, only a fraction of telomeres are still enriched with 8oxoG when replication forks traverse them. To address this, and further confirm the genome instability we’ve observed is due to replication stress, we synchronized cells at the G1/S boundary or mid S-phase prior to telomeric 8oxoG induction (Figure 4 and Figure S4). Compared to the asynchronous control, 20 hour thymidine block did not significantly raise the background level of MN. However, after 8oxoG induction and 0.05 μM VE822, both U2OS and HeLa cells displayed significantly more MN when damaged at G1/S, and especially mid S-phase compared to the asynchronous control.

**Figure 4:**
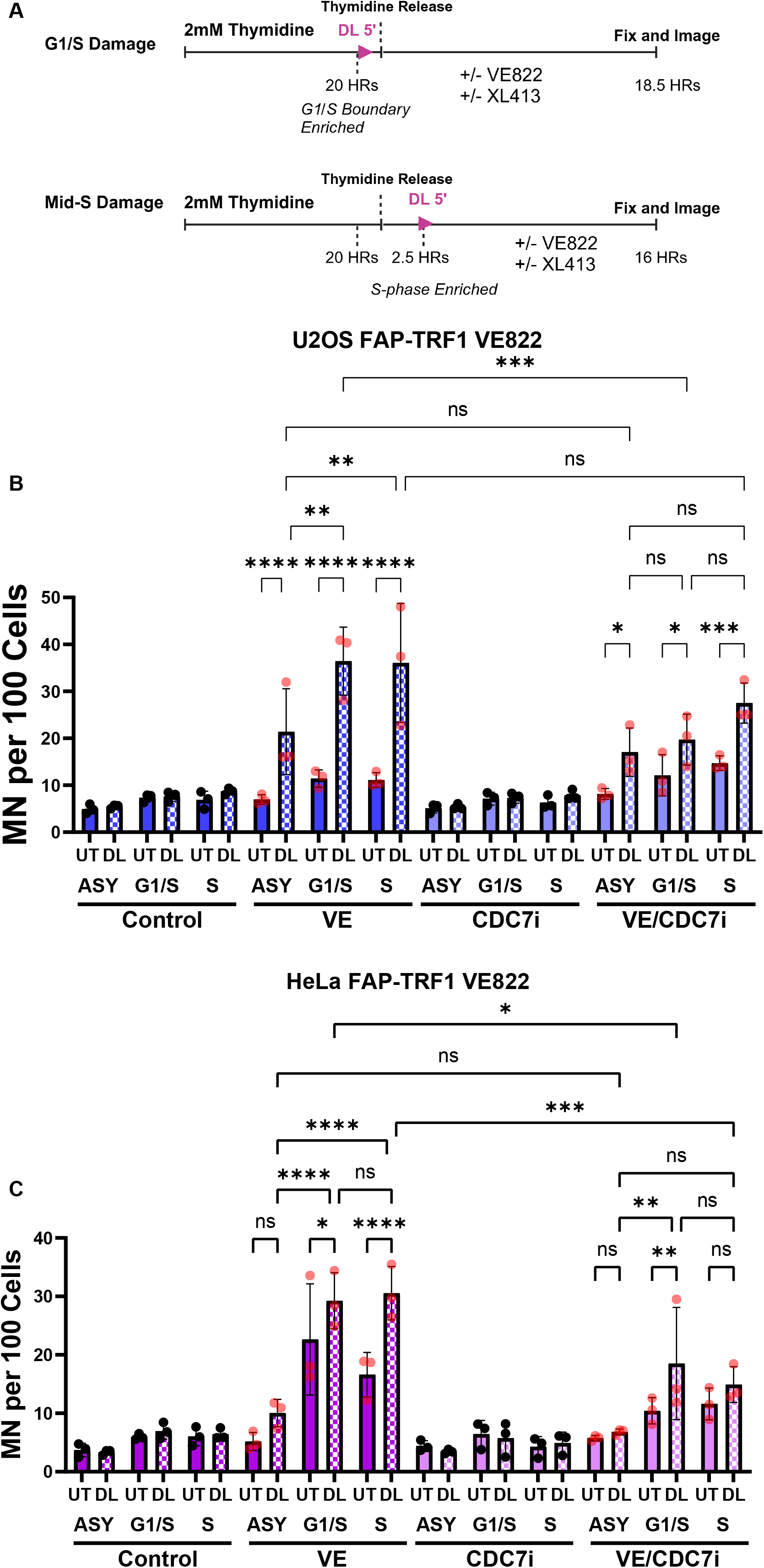
DNA Replication is Hypersensitive to Telomeric 8oxoG. (A) Schematic of cell synchronization experiments. Cells were arrested at G1/S boundary with thymidine for 20 hours. G1/S damaged cells were not released prior to dye and light. Mid-S damaged cells were released 2.5 hours before dye and light treatment. Drugs were added after dye and light in both cases. U2OS (B) and HeLa (C) FAP-TRF1 cells were treated as indicated (XL413 CDC7i = 0.1 μM U2OS and 2 μM HeLa) and MN scored 18.5 hrs after thymidine release. All data are mean ± SD. Each point is a biological replicate. All data analyzed by 2-way ANOVA.

### CDC7 Inhibition Mitigates Telomere Replication Stress

The potential reasons for ATRi induced MN following telomere replication stress are numerous. Previous studies have shown ATRi can induce dormant origin firing, replication fork degradation, nucleotide depletion, and premature mitotic entry^23^. CDC7 was shown to induce dormant origin firing when ATR is inhibited, and also promote DSBs and DSB signaling in hydroxyurea and ATRi treated cells^24, 25^. Given these two critical roles of CDC7 during replication stress, we treated synchronized cells with a well characterized CDC7i (XL413) alongside VE822. We used different doses of XL413 for HeLa and U2OS, as we found doses above 2 μM for HeLa or 0.1 μM for U2OS significantly raised background MN in our experiments (data not shown). Using the selected doses, CDC7i alone had no effect on MN in either cell line. However, when XL413 was added with VE822, we saw a significant reduction in MN compared to VE822 alone. Dormant origin firing within a telomere was previously observed in cells treated with aphidicolin^26^.

### Premature Mitotic Entry Induces MN In Cells with Telomeric 8oxoG

While telomeric 8oxoG reduces S-phase (EdU+) wild-type RPE cells, it does not change the percent of U2OS, HeLa, or p53ko RPE in S-phase (Figure 5A). ATRi alone reduces EdU+ cells, but there is no further change with dye and light for this phenotype. This suggests that perhaps ATRi is affecting the G2/M phase checkpoint more than the intra-S phase checkpoint in our system (Figure 5B). Consistent with this, when cells are treated with a Wee1 inhibitor after 8oxoG induction at telomeres, we observed a dose-dependent increase in MN (Figure 5C, D). Although Wee1 can affect S-phase^27^, its most understood role is controlling CDK1 phosphorylation to regulate the G2 to M-phase transition. Aside from regulating replication forks directly, ATR and Chk1 regulate CDC25 phosphatases which also control CDK1. This suggests the mechanism for ATRi induced MN following telomere replication stress is due to premature mitotic entry.

**Figure 5:**
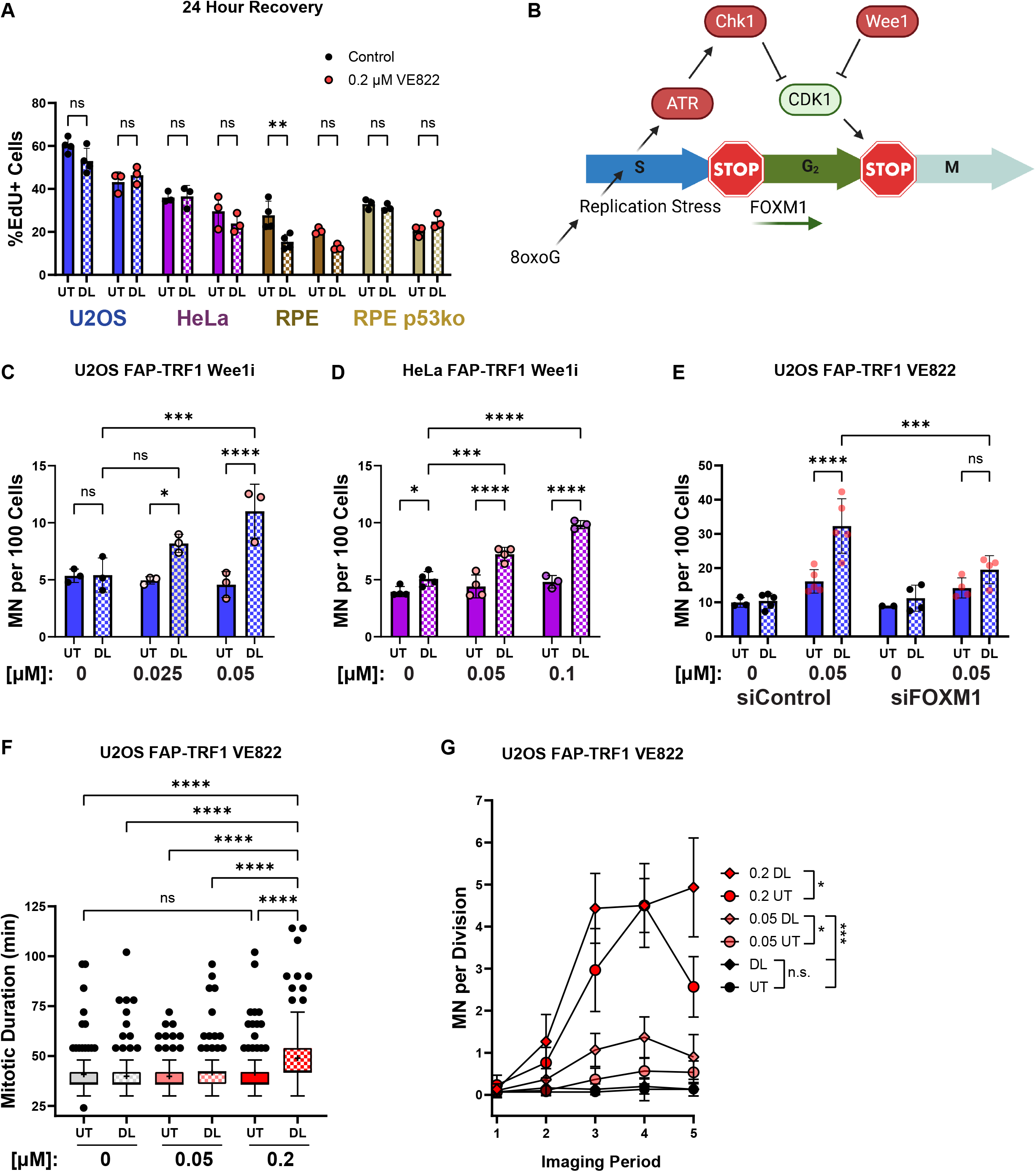
Premature Mitotic Entry Induces MN In Cells with Telomeric 8oxoG. (A) 1 hour before fixation, cells were pulsed with EdU. Fixation was 24 hours after treatment with dye and light for 5 minutes. The percent of EdU positive cells was scored by microscopy. (B) Schematic of ATR/Chk1, Wee1, and FOXM1 roles in G2 and G2/M checkpoint. (C, D) U2OS and HeLa FAP-TRF1 cells were treated as in A, but recovered with Wee1i MK-1775. (E) U2OS FAP-TRF1 cells were treated with control or FOXM1 siRNA for 48 hrs before treating with dye and light for 5 minutes and recovered 24 hours with or without VE822. Data are mean ± SD. Each point is a biological replicate. (F) U2OS cells expressing H2B-mCherry were treated as indicated, and imaged for 36 hours on a stage-top incubator. For each mitotic event scored, the duration of mitosis (see methods) was quantified. Data are Tukey box-plots, with “+” representing the mean. N=150 mitotic events for each condition. (G) The mitotic events from (F) were scored for the number of MN produced. Each condition was divided into 5 windows of analysis (1 = 2-8.8, 2 = 8.8-15.6, 3= 15.6-22.4, 4= 22.4-29.2, 5= 29.2-36 hrs). Data are the mean with 95% confidence intervals. N=30 per time period per condition. All data analyzed by 2-way ANOVA. In G, post-hoc was restricted to comparing conditions, not times.

Previous studies have shown ATRi induces the transcription of G2/M-phase related genes in a FOXM1 dependent fashion^28^. We confirmed in our system, that knockdown of FOXM1 abrogated the genome instability observed after ATRi and telomere replication stress (Figure 5E). This is consistent with a mechanism that when the RSR is inhibited following telomere replication stress, cells prematurely exit G2 and enter mitosis.

A phenotype of cells attempting to divide with unresolved replication intermediates is prolonged mitosis^29^. Mitosis is such a critical aspect of the cell cycle that its timing is conserved throughout different cell types in humans, and even small changes in duration can signal cell death or growth arrest. We labeled chromatin in U2OS FAP-TRF1 cells with H2B-RFP and scored mitotic events after induction of telomeric 8oxoG and ATRi. We divided the experiment into 5 windows (Imaging Periods) to ensure an even sampling of events. For each event we calculated the length of mitosis from nuclear envelope breakdown (NEBD) to completion of cytokinesis (Figure S5 and methods). Cells treated with dye and light and 0.2 μM VE822 showed a modest, but statistically significant increase in miotic duration, and were the only cells with a mitotic duration significantly different from all other conditions (Figure 5F).

We evaluated the number of MN produced from each mitosis scored throughout the imaging experiment (Figure 5G). Consistent with the effects of 8oxoG requiring an S-phase to raise MN levels, no condition showed increased MN during the first imaging period (up to ∼8.8 hrs post treatment). An increase in the number of MN per mitosis began in the second imaging period, and by the third period (up to 22.4 hrs post treatment) the average MN per division for dye and light and 0.2 μM VE822 treated cells was ∼4, and reached ∼5 by period 5. Except for period 4, 0.2 μM VE822 dye and light treated cells had more MN per division than 0.2 μM VE822 UT cells, and overall was significantly different. 0.05 μM VE822 dye and light treated cells showed significantly more MN than 0.05 μM VE822 UT, averaging ∼1 MN per division in periods 3-5, dramatically higher than the background rate of ∼0.1 MN per division for the controls. 0.05 μM UT never averaged > 0.6 MN per division, and was not significantly different from the no drug controls. In summary, ATRi and oxidative telomere damage synergize to disrupt mitosis, resulting in a longer mitosis and more MN per division.

### Extended G2, but not Mitosis, Rescues Genome Instability after Oxidative Telomere Damage

To directly test if restraining entry into mitosis would allow cells time to recover from telomere 8oxoG and RSR inhibition, we used the CDK1 inhibitor RO-3306 to control mitotic entry (Figure 6A). When asynchronous U2OS and RPE cells were treated with CDK1i alongside 0.2 μM VE822, this completely prevented MN induction as cells were unable to leave G2-phase (Figure 6B, C). To determine if a temporary hold in G2 could rescue ATRi induced MN after oxidative telomere damage, we treated cells with dye and light as before, and the incubated with VE822 and/or CDK1i for 16 hours. Then these cells were “released” from drug, and provided fresh media with no drug for 8 hours. Cells treated only with VE822 for 16 hours still had a significant increase in MN, albeit less than those treated for 24 hours. However, cells co-treated with VE822 and CDK1i for 16 hours showed a dramatic reduction in MN, significantly less than the VE822 singly treated cells and not significantly greater than the no drug control. This strongly suggests the role of ATR signaling following telomere replication stress is prevention of premature mitotic entry.

**Figure 6:**
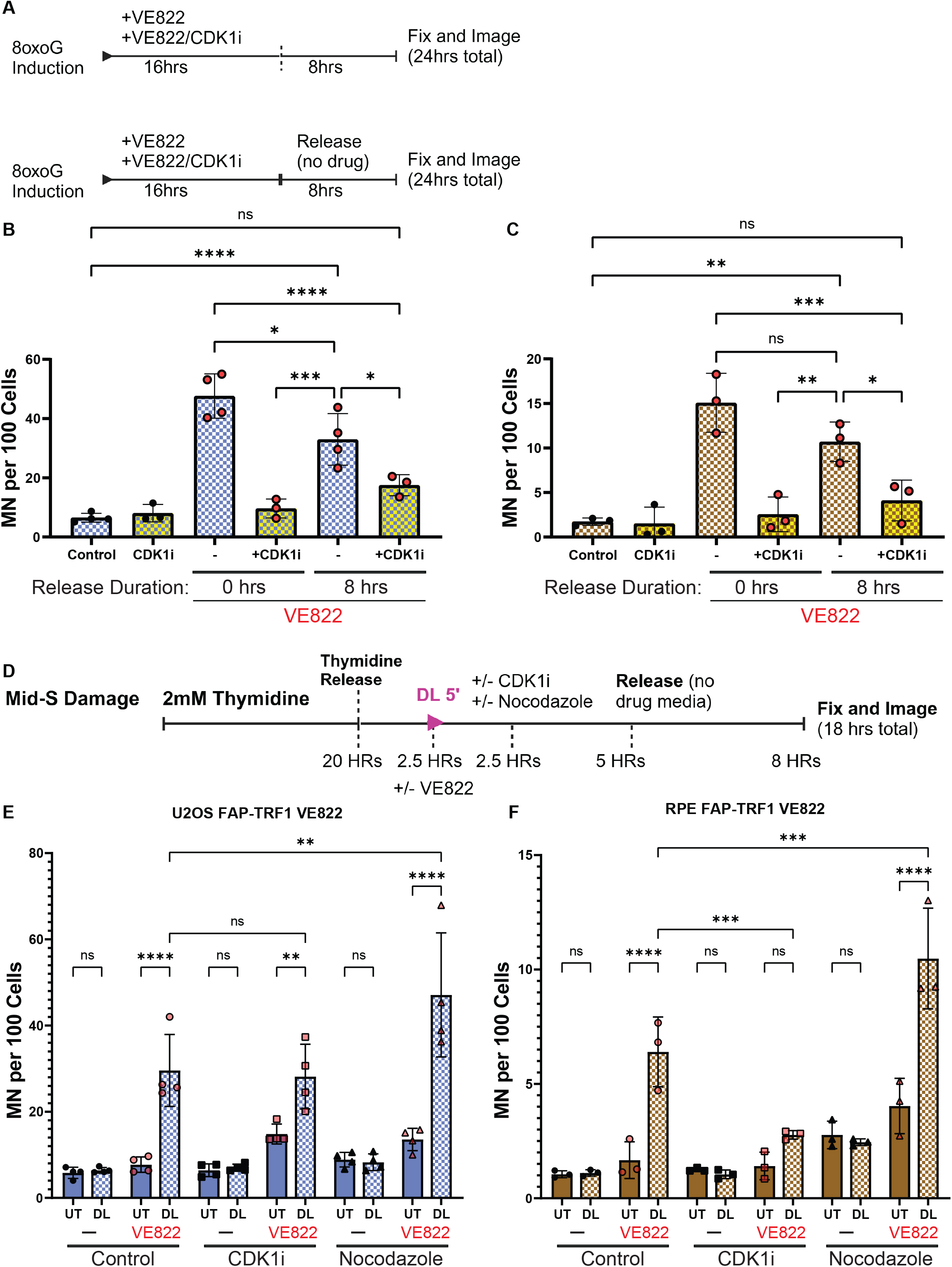
Extended G2, but not Mitosis, Rescues Genome Instability after Oxidative Telomere Damage. (A) Schematic of CDK1i release experiment. After 5 min dye and light treatment, cells were treated with 0.2 μM VE822 or 10 μM RO-3306, and recovered 16 or 24 hours. Cells recovered 16 hours, were washed and released into drug free media for 8 hours. U2OS (B) and RPE FAP-TRF1 cells (C) were treated as indicated and in (A) and MN scored. (D) Schematic for the CDK1 and nocodazole synchronization experiment. Cells were treated with dye and light for 5 mins in Mid-S-phase. 0.05 (U2OS) or 0.2 μM VE822 (RPE) was added immediately after 8oxoG induction. 2.5 hrs later, 10 μM RO-3306 or 50 ng/ml nocodazole were added for another 5 hours, before washing and release into no drug media. U2OS (E) and RPE FAP-TRF1 cells (F) were treated as indicated and in (D) and MN scored. All data are mean ± SD. Each point is a biological replicate. All data analyzed by 2-way ANOVA.

Work by us and other groups have shown DNA repair synthesis in mitosis (MiDAS) is activated after telomere replication stress, and this is presumed to be cytoprotective^5^. We wondered if a brief mitotic arrest would similarly protect ATRi treated cells following telomere oxidative damage, as observed with G2 arrest. To accommodate the fact prolonged mitotic arrest increases genome and specifically, telomere instability, we conducted cell synchronization^30^. Cells were only treated with 50 ng/ml nocodazole for 5 hours to maintain low MN background, as holds for 10-24 hours caused massive genome instability in U2OS cells (data not shown). U2OS and RPE FAP-TRF1 cells were treated to induce telomeric 8oxoG in mid S-phase and after 2.5 hour recovery, were treated with CDK1i to prevent G2-phase exit, or nocodazole to prevent M-phase exit. 5 hours later, cells were released from the hold, and given 8 hours to complete division without any drug present. With the chosen conditions, nocodazole alone did not raise MN levels compared to the UT control, and only modestly increased MN in RPE cells co-treated with VE822. However, when cells in S-phase were treated with dye and light and ATRi, nocodazole significantly increased MN compared to cells treated with only dye and light and ATRi. Under these conditions CDK1i did not rescue MN for U2OS cells (but also did not raise MN), perhaps due to the synchronization, but did rescue RPE cells. In total, our data suggest allowing cells increased time in G2-phase, but not M-phase, following telomere 8oxoG induction and ATR inhibition allows them to resolve replication stress intermediates that when transmitted to mitosis, cause serious genomic instability.

## Discussion

Selective perturbation of cancer cells over non-diseased tissue is paramount for comprehensive cancer care. Tremendous advancements have been made using DNA damaging treatments such as platinum, nucleotide analogues, and ionizing radiation, however they lack an ability to selectively impact cancer cells. More targeted therapies using inhibitors of the DNA damage response such as PARP inhibitors have gained significant use in homologous recombination deficient tumors because they exploit a vulnerability specific to the cancer, elevating their sensitivity to DNA damage.

Here, we find that by inducing replication stress at telomeres through oxidative DNA damage, we can sensitize cancer cells to low dose ATR and Chk1 inhibition. We validated our findings using two ATR and Chk1 inhibitors at different doses, in 4 cancer cell lines. Additionally, we used two lines each, that maintain their telomeres with telomerase, or the ALT pathway to ensure there was no dependence on telomere maintenance mechanism (Figure 1). Coincident with the increase in genome instability, we found reduced cancer cell viability and persistent telomere dysfunction after 8oxoG induction and ATRi, suggesting this treatment paradigm could be effective in stunting cancer growth in patients (Figure 2).

Importantly, while non-cancerous, otherwise normal cell lines senesce following telomeric 8oxoG induction, they were unaffected further by low doses of ATRi (Figure 3). This presents a possible therapeutic window in which targeted replication stress induction in cancer cells, in combination with low dose ATRi, could cancer cell death, while leaving normal tissue unharmed. We hypothesized the reason normal cells were less affected by this combination treatment was due to intact p53 signaling. And indeed, knockout of p53 significantly sensitized these cells indicating a cytoprotective function of p53 in the context of telomere replication stress and ATR inhibition. Interestingly, 3 of the cancer lines we studied have dysfunctional p53 signaling. The only exception being the LOX melanoma line, which also had the weakest response to ATRi and dye and light treatment^31^. We speculate that p53 may have a similar importance in modulating resistance to telomere replication stress in cancer cells, but is just more often than not dysregulated.

To understand mechanistically why ATRi caused genome instability after 8oxoG formation in telomeres, as manifested by increased MN, we turned to synchronization. Previously we showed the DDR at telomeres after 8oxoG induction was blunted in serum-starved cells, which could not enter S-phase, demonstrating the effects of telomeric 8oxoG observed were due to perturbed DNA replication^20^. Here, we enriched cells in G1/S and S, and found telomeric 8oxoG and ATRi induced more MN than in asynchronous cells. This confirms the MN we have found in our study arise from replication stress, as enriching the replicating population increased the phenotype.

ATR inhibition has been shown to induce dormant origin firing, prevent recruitment of repair factors to stalled forks, and disrupt cell cycle progression. In our system, while ATRi reduces the number of S-phase cells 24 hours after treatment, telomeric 8oxoG does not impart this specific phenotype. Instead, we suspected ATRi allows cells to progress into mitosis prematurely, before resolving intermediates or under-replicated DNA arising from telomere replication stress. Supporting this, promoting mitotic entry with Wee1 inhibition also increased MN after oxidative damage to telomeres, and the increase in MN after ATRi could be rescued by FOXM1 knockdown. FOXM1 promotes premature mitotic entry in cells treated with ATRi or Chk1i, so we concluded that ATRi allowed cells with unresolved telomere replication intermediates to enter mitosis, which lead to chromosomal instability^32^.

Consistent with the notion ATRi allowed cells to enter mitosis prematurely with unresolved telomere damage, we observed an increase in the length of mitosis in cells after 8oxoG induction and 0.2 μM VE822. Several recent studies have shown prolonged mitosis after replication stress and this correlates with increased genome instability and or cell death and arrest^33-35^. Furthermore, earlier work has shown a telomere-specific cohesion defect can also lead to prolonged mitosis^11, 36^. While we have not assessed cohesion status in our system, it is tempting to speculate the difficulties caused by telomeric 8oxoG may impair sister telomere separation in mitosis.

We surmised oxidative base damage to telomeres produces replication intermediates that when transmitted to mitosis due to ATRi, activate the spindle assembly checkpoint, delaying progression. We tested this hypothesis by blocking premature G2-phase exit with a CDK1i, before release into mitosis. Impressively, we found this delay in mitotic entry in the face of ATRi rescued the genome instability phenotype. Moreover, we confirmed this was due to a G2 specific hold, and not a division hold, by comparing CDK1i with nocodazole. Even though we optimized conditions so that VE822 and nocodazole alone or in combination did not increase MN, we saw a significant increase in MN in cells with telomeric 8oxoG treated with both drugs. This shows not only that prolonging mitosis after telomere damage does not allow for repair, but that it actually exacerbates the problem.

In conclusion, we have determined targeted oxidative damage to telomeres can sensitize cancer cells to low dose ATR signaling inhibition. This has important relevance for cancer treatment, because the conditions used here show non-cancerous cells are resistant to low dose ATRi, which may translate to a higher tolerance for patients. These are the principal coordinators of the replication stress response in human cells, and are currently being pursued by several companies for mono- and combination cancer therapies. Therefore, it is critical we continue to develop treatment paradigms that will sensitize cancer cells to the inhibitors, at doses which do not affect non-diseased cells. We predict that at worst, non-cancer cells will senesce under these conditions, leaving them benign or mildly immunogenic.

## Figure Legends

Figure S1: (A) 4 day cell growth assay. Cells were treated with the indicated doses of VE822 and counted 4 days later. Data normalized to no VE822 control. (B) Representative DAPI staining of U2OS cells with micronuclei. (E, F) SAOS and LOX FAP-TRF1 cells were treated with dye and light for 5 minutes and MN counted 24 hours later. (G, H) U2OS and HeLa FAP-TRF1 cells were treated with dye and light for 5 minutes and recovered 24 hours with 0.2 μM VE822 or 1.0 μM ATMi (KU60019). (I, J) as in (G, H) except cells recovered with Chk1/2 inhibitor AZD7762.

Figure S2: Representative scatter-plot of U2OS FAP-TRF1 flow cytometry gating.

Figure S3: (A) RPE FAP-TRF1 colony survival after 5 minute dye and light treatment and recovered for 7-10 days with AZD6738. (B) 24 hours after 5 minute dye and light treatment, RPE FAP-TRF1 cells were scored for MN. (C) As in Fig 2, RPE cell apoptosis and necrosis measured 4 days post treatment. (D) Wild-type and p53ko BJ FAP-TRF1 cells were treated with dye and light for 5 minutes, and recovered for 4 days before counting. Data are normalized to the UT no drug for each cell line. (E) 24 hours after treatment, cells were scored for MN. (F) 24 hours after treatment, LM216J FAP-TRF1 cells were scored for MN. All data are mean ± SD. Each point is a biological replicate. All data analyzed by 2-way ANOVA.

Figure S4: Representative propidium iodine flow cytometry of RPE (A) and U2OS (B) FAP-TRF1 cells for the experiments in Figure 4 and Figure 6.

Figure S5: (A) Immunoblot showing FOXM1 knockdown. (B) Representative mitosis from U2OS FAP-TRF1 cells showing H2B-RFP labeled chromatin and DIC imaging of the cell. Key mitotic events are noted as well as relative time. (C) Data from Figure 5 F and G, plotted to show each mitotic duration and the number of MN produced throughout the experiment.

## Methods

### Cell Culture

FAP-mCer-TRF1 HeLa, U2OS, BJ hTERT, and LM216J cells were cultured in DMEM, RPE hTERT in DMEM/F12, SAOS in McCoy’s 5A, and LOX IMVI in RPMI all with 10% FBS, 1% penicillin/streptomycin, and 500 μg/ml G418 to maintain FAP-mCer-TRF1. BJ cells received Cytiva Hyclone FBS while all others received Gibco Value FBS. All cells were cultured at 5% CO_2_ and O_2_ at 37°C. All lines were generated in previous publications^18-20^.

### Cell treatments

For acute treatments with MG2I dye and light, cells were pre-treated with Opti-MEM (GIBCO) for 15 min at 37°C, followed by treatment with 100 nM MG2I dye for 15 min. Cells were then exposed to a high intensity 660 nm LED light at 153 mW cm^-2^, as previously described. For a 5 minute treatment, this results in ∼46 J cm^-2^ of energy. Lightbox specifications were as described previously, except the glass stage was replaced with a holder for a culture flask or multi-well dish, the light path was 11 cm, and another fan was added for heat dissipation^37^. MG2I was provided by Brigitte F. Schmidt at Carnegie Mellon University.

ATR, ATM, CDC7, Chk1, CDK1, and WEE1 inhibitors were added after dye and light treatment unless otherwise indicated. Expect for CDC7i XL413 which was PBS, all drugs were dissolved in DMSO at 10 mM, and diluted in media for treatments from aliquots.

### Cell Growth

Cells were plated in 6-well plates at 2×10^4^ per well. Cells were treated with dye and light the next day, and drugs added after. Cells recovered 4 days with drug, and then were counted with Denovix Cell Drop. Each condition had technical duplicate or triplicates.

### Colony Survival

Cells were plated at low density (1-2×10^2^). Cells were treated with dye and light the next day, and drugs added after. Cells recovered 7-10 days with drug, and then were fixed with ice-cold methanol on ice, and stained with 0.1% crystal violet. After washing with water and drying, colonies were manually counted. Each condition had technical duplicates or triplicates.

### Micronuclei Quantification

Approximately 1×10^5^ cells were seeded on 22×22 mm cover glass. Next day cells were treated as indicated and typically recovered for 24 hours with drug. Cells were fixed with 4% formaldehyde diluted in PBS and stained with DAPI diluted in water before mounting. Each sample was imaged at 40x at 10-25 fields of view depending on cell density on a Ti2 Eclipse widefield microscope (Nikon) equipped with a Spectra III Light Engine (Lumencore) and ORCA-Fusion BT Digital sCMOS camera (Hamamatsu). Cells were counted automatically with Nikon Elements Bio Analysis Cell Count Module, and micronuclei were manually counted. Each sample had 300-1000 cells counted.

### Immunofluorescence Microscopy and FISH

Cells were seeded and treated similarly for the MN quantification. For RPA staining, cells were pre-treated with CSK extraction buffer on ice (100 mM NaCl, 3 mM MgCl2, 300 mM glucose, 10 mM Pipes pH 6.8, 0.5% Triton X-100), before fixing. Cells were fixed on ice in cold methanol, and blocked with 10% normal goat serum, 1% BSA and 0.1% Triton X in PBS, 1 hour RT. Cells were incubated overnight at 4°C with primary antibodies. The next day, cells were washed three times with PBS-T before incubating with secondary antibodies and washing again three times with PBS-T. For FISH, the cells were refixed with 4% formaldehyde, rinsed with 1% BSA in PBS and then dehydrated with 70%, 90% and 100% ethanol for 5□min. Telomeric PNA probe was diluted 1:100 (PNABio) prepared in 70% formamide, 10 mM Tris-HCl pH 7.5, 1× Maleic Acid buffer, 1× MgCl2 buffer and boiled for 5 min before returning to ice. Coverslips were then hybridized in humid chambers at room temperature for 2 h or overnight at 4°C. The cells were washed twice with 70% formamide and 10 mM Tris-HCl pH 7.5, three times with PBS-T then rinsed in water before staining with 4′,6-diamidino-2-phenylindole (DAPI) and mounting. Images were acquired with a 60x oil objective, and deconvolved before analysis with Nikon AR.

### Live-Cell Imaging

Cells were seeded in glass bottom 6-well plates and recovered overnight. After treatment, cells were swapped to phenol red-free DMEM supplemented with 10% FBS with or without ATR inhibitor for imaging and placed onto the preincubated microscope stage. Images were acquired every 6 minutes for 36 hours using a RFP filter set (20 millisecond exposure, 5% laser power) and differential interference contrast channels (DIC, 100ms exposure). 2×2 images were stitched using NIS Elements, with a 2% overlap stitching performed in the RFP channel.

Raw images were processed using a custom filter to identify mitotic cells in an unbiased manner. This filter identified rounded cells above a diameter threshold by identification of high-contrast circular spots in the DIC channel. The first two hours of imaging were discarded to allow cells to adjust to the stage-top incubator. The remaining 34 hours were split into five equal periods of 408 minutes each. The first ten in-focus cells from the five time periods that were identified by the mitotic filter were selected for analysis to ensure even sampling across the entire imaging duration. Identified cells were then analyzed manually with the following events recorded: Nuclear envelope breakdown (NEBD), metaphase, anaphase, G1 begin. These events were defined: NEBD – condensation of the nucleus visualized with H2B-RFP; Metaphase-bright vertical alignment of H2B-RFP signal; Anaphase-first frame of separation of the H2B-RFP signal from the vertical alignment seen in metaphase in previous frames; G1 begin-completion of cytokinesis indicated by a relaxation of the plasma membrane or diffusion of RFP signal. The timepoint these events occurred was recorded for each cell in each experimental condition and transformed into event duration times, with reported values from the 50 analyzed cells pooled. The number of MN produced from each division was also recorded. Three biological replicates were analyzed for each indicated experimental condition.

### Immunoblotting

Cells were collected by scraping on ice and washed with PBS. Each pellet was lysed on ice with 1x RIPA buffer supplemented with 1x Halt Protease and Phosphatase Inhibitors, 1 nM PMSF, and Benzonase for 15 minutes and 37°C for 10 minutes. After centrifugation, protein concentrations were determined with BCA assay and 15-50 μg was electrophoresed on Bis-Tris gels. Protein was transferred to PDVF membranes, blocked with 5% milk and incubated with primary antibody overnight. Licor fluorescent secondary antibodies were applied next day and signal detected on Licor Odyssey.

### Flow Cytometry

Cell cycle: Cells were trypsinized and washed with PBS before fixing with ice-cold 70% ethanol for at least 3 hours. After fixation, cells were washed once with PBS and resuspended in FxCycle™ PI/RNase Staining Solution (Invitrogen) and incubated for 15-30 minutes according to the manufacturer’s protocol. Cells were analyzed using an Attune NxT acoustic focusing flow cytometer (Thermo Fisher Scientific). Gating was performed using FlowJo (BD Biosciences).

Cell death: 2×10^5^ cells were plated in 60 mm dishes. The next day, cells were treated with dye and light, followed by the addition of ATR inhibitor. After 96 hours, supernatant was collected, and cells were washed with PBS, detached with trypsin, and centrifugated. Cell pellets were resuspended in 100 μL of 1X binding buffer (Invitrogen). 10 μL of the Annexin V, fluorescein conjugate (FITC annexin V) (Invitrogen), and 10 μL of the propidium iodide with RNAse (50 μg/mL) were added, and cells were incubated in the dark for 15 min at room temperature. After the incubation, 400 μL of binding buffer was added. Cells were acquired using Attune NxT acoustic focusing cytometer (Thermo Fisher Scientific) and analyzed with FlowJo (BD Biosciences).

## Author Contributions

AG conducted and analyzed most experiments. NCM performed apoptosis and immunoblotting experiments. AD conducted the IF/FISH experiments. PLO supervised, and contributed to writing and editing the manuscript. RPB conducted experiments, designed the study and data presentation, and wrote the manuscript.

## Acknowledgements

We thank Anna Holland with helping image the ATMi and Wee1i experiment, and others. We acknowledge the Flow Cytometry Core Laboratory, which is sponsored, in part, by the NIH/NIGMS COBRE grant P30 GM103326 and the NIH/NCI Cancer Center grant P30 CA168524. We thank Dr. Brigette Schmidt at Carnegie Mellon University for MG2I dye. This work was supported by R35ES030396 to PLO and R00ES033771 and KUMC Start-up Funding to RPB.

