## Supplementary figures and images for "Oxidative Base Damage to Telomeres Sensitizes Cancer Cells to ATR Inhibition"

### Supplemental Figure 1

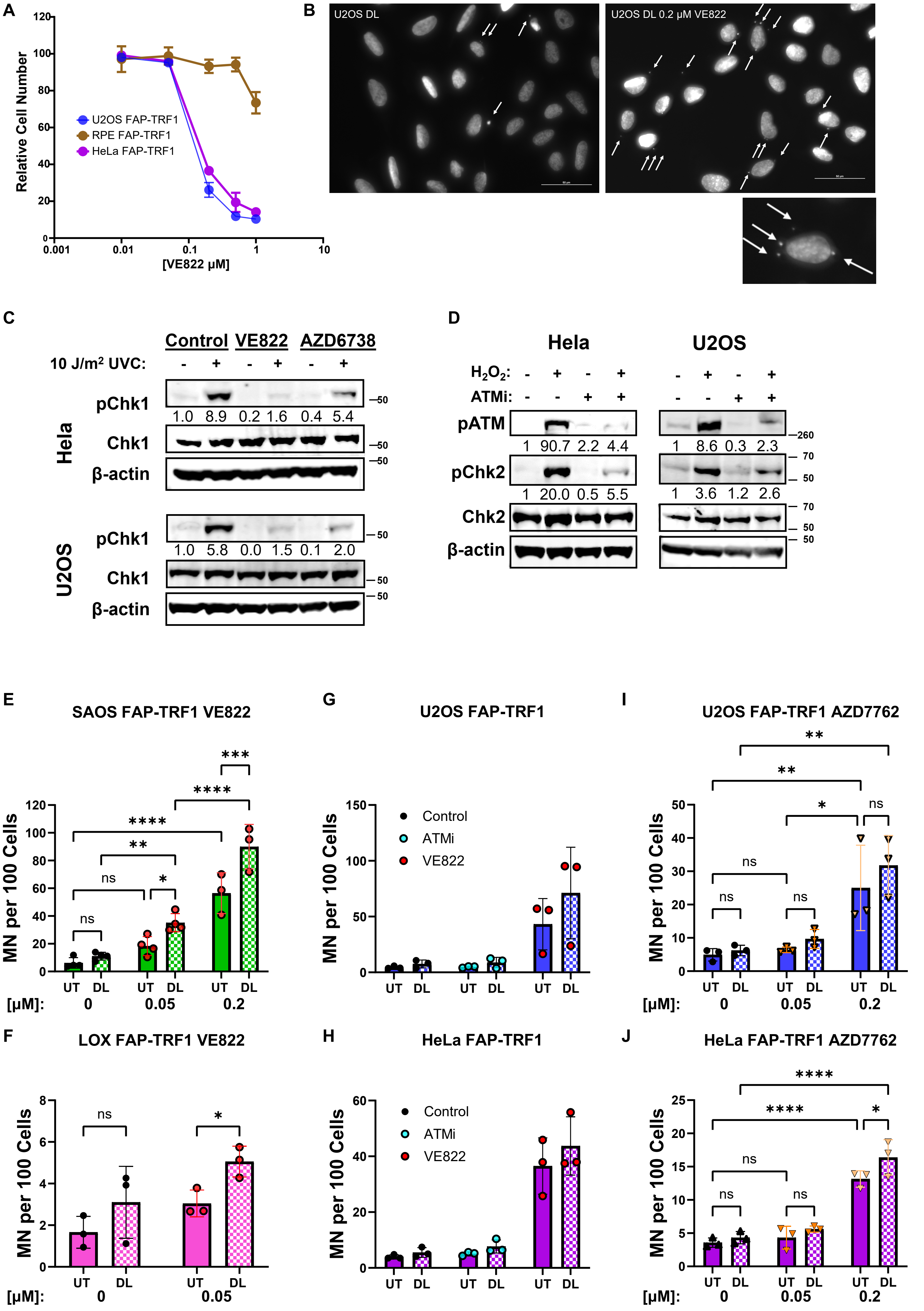

### Supplemental Figure 2

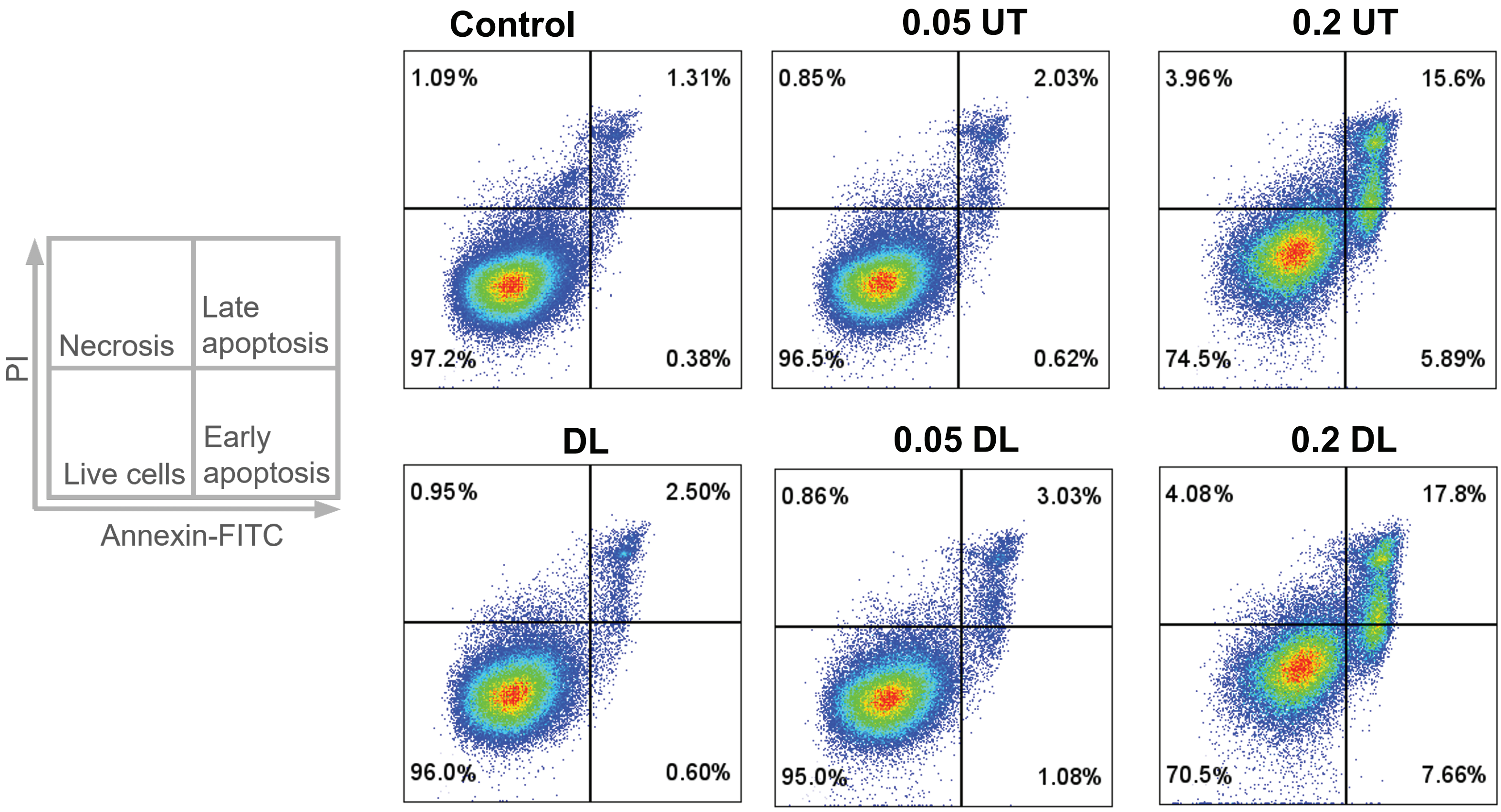

### Supplemental Figure 3

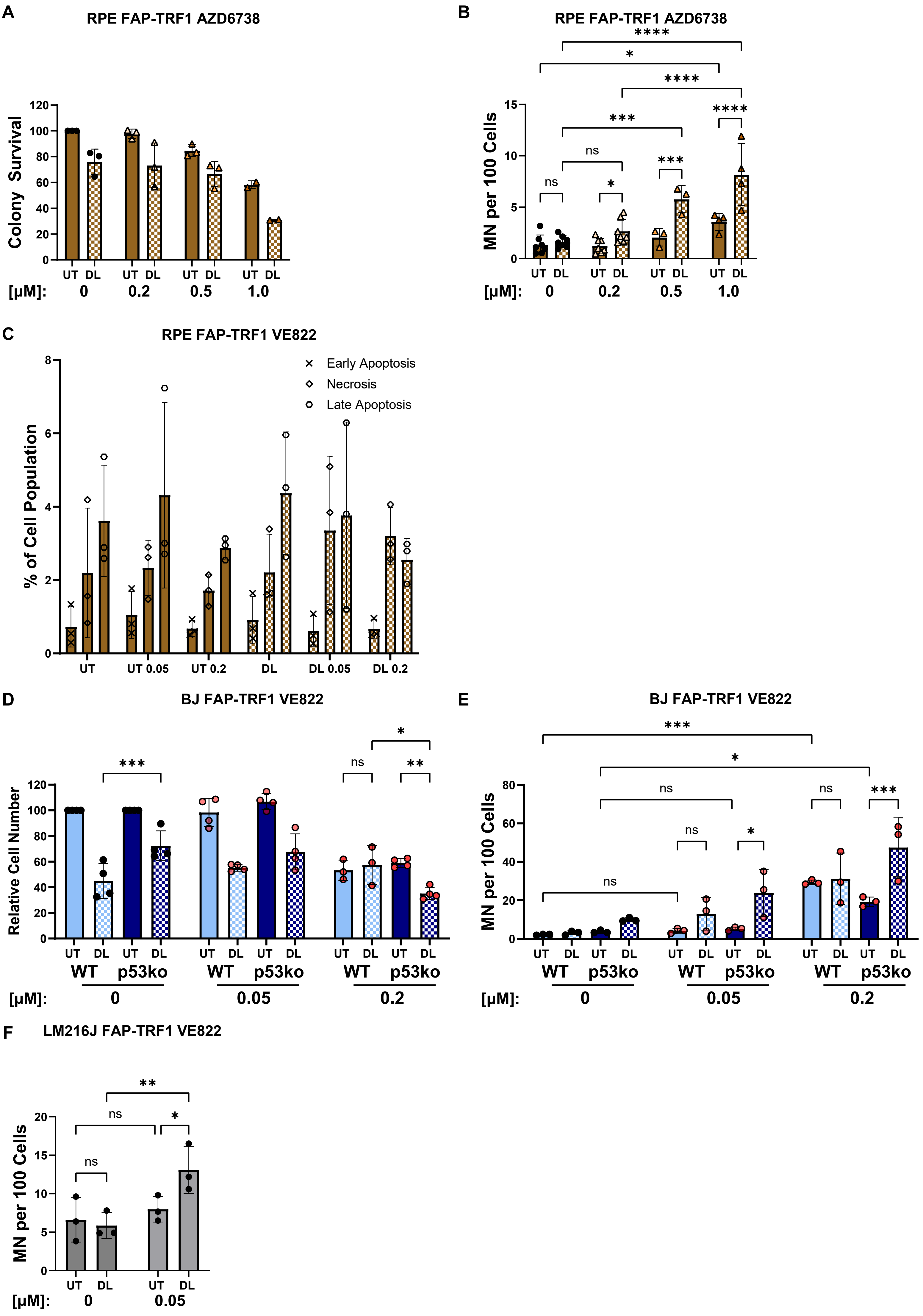

### Supplemental Figure 4a

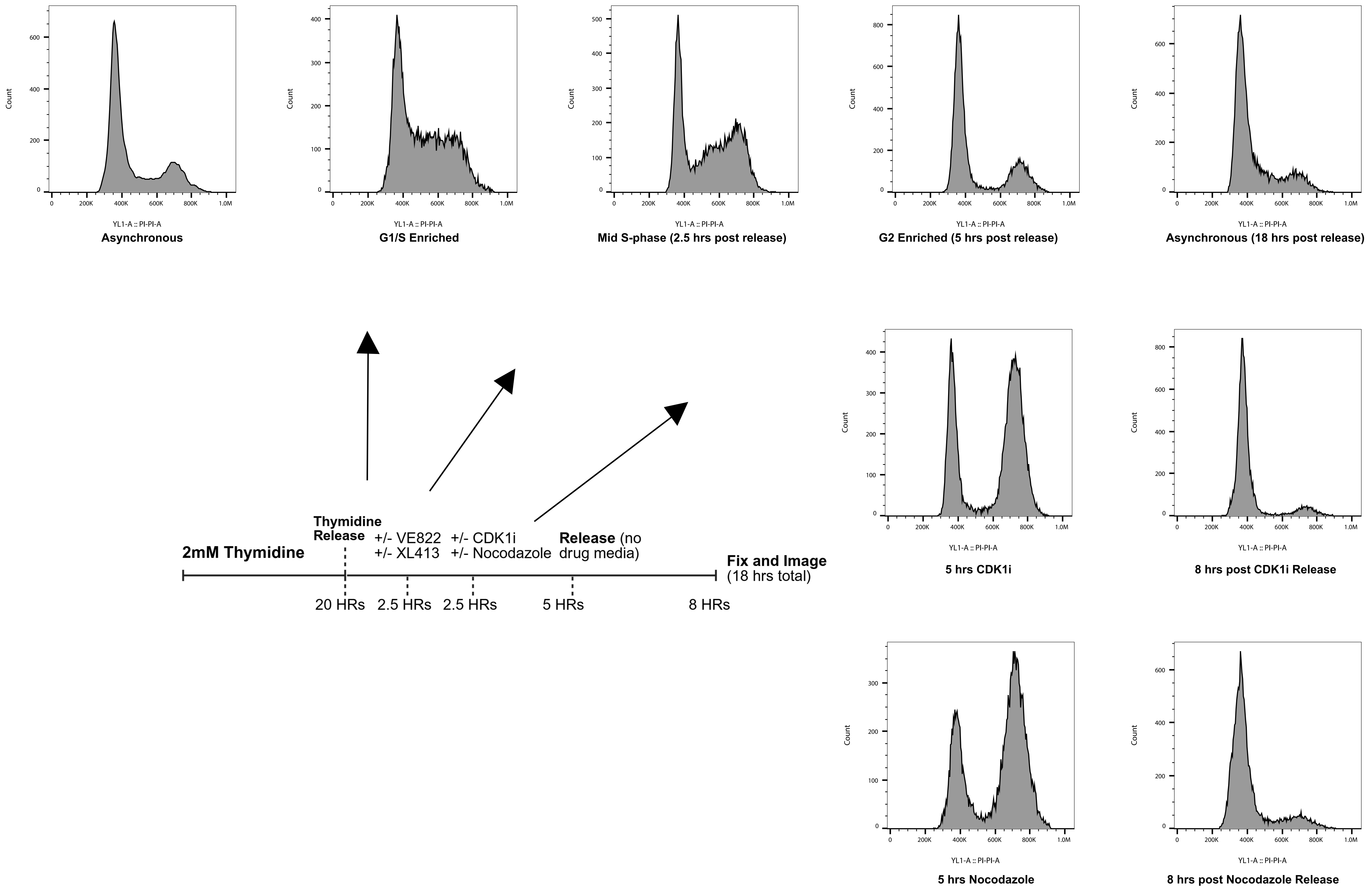

### Supplemental Figure 4b

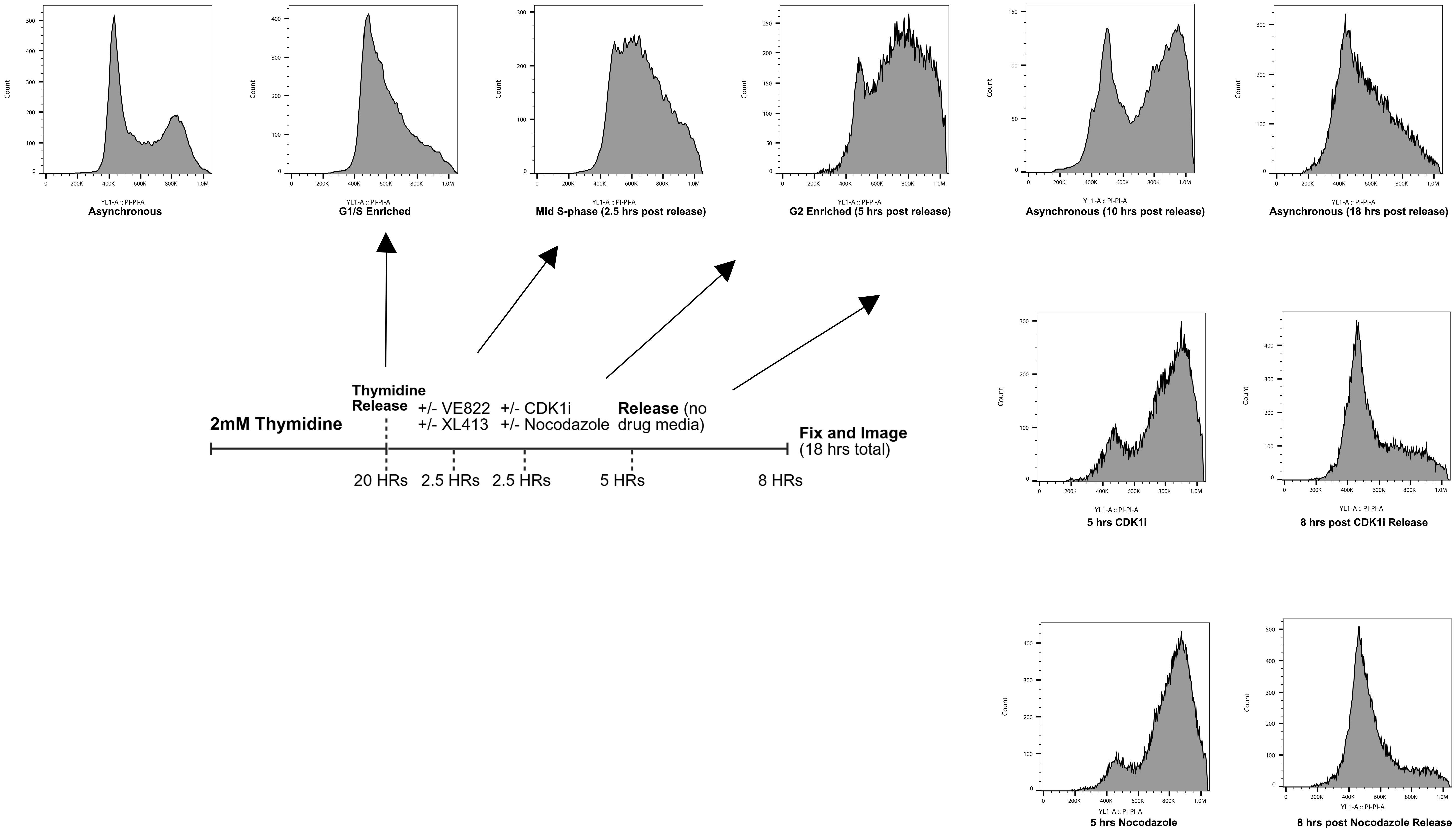

### Supplemental Figure 5

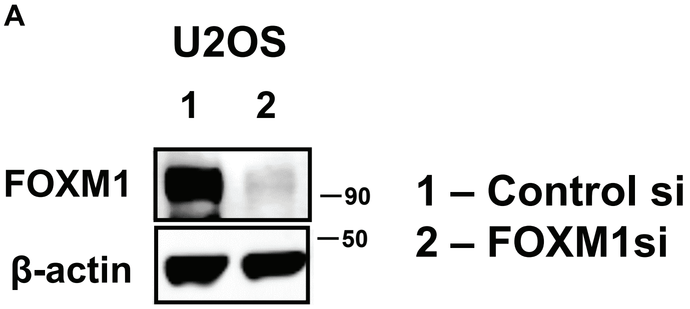

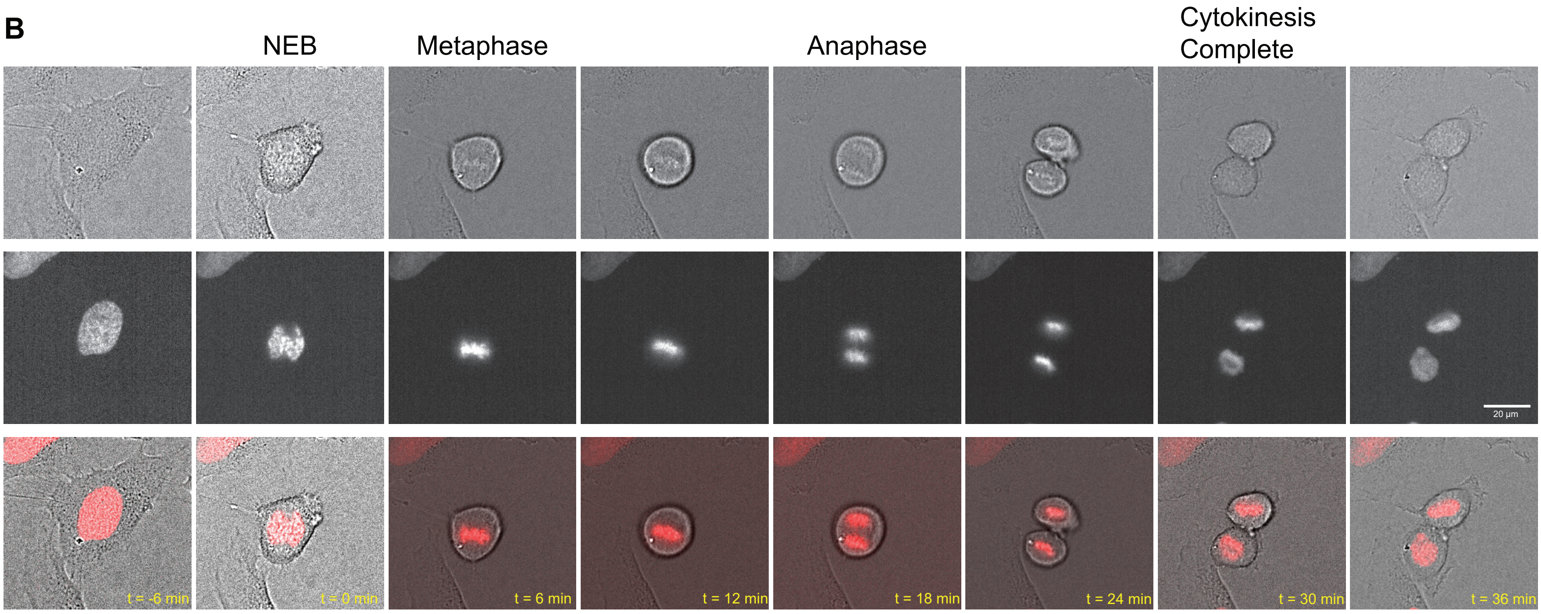

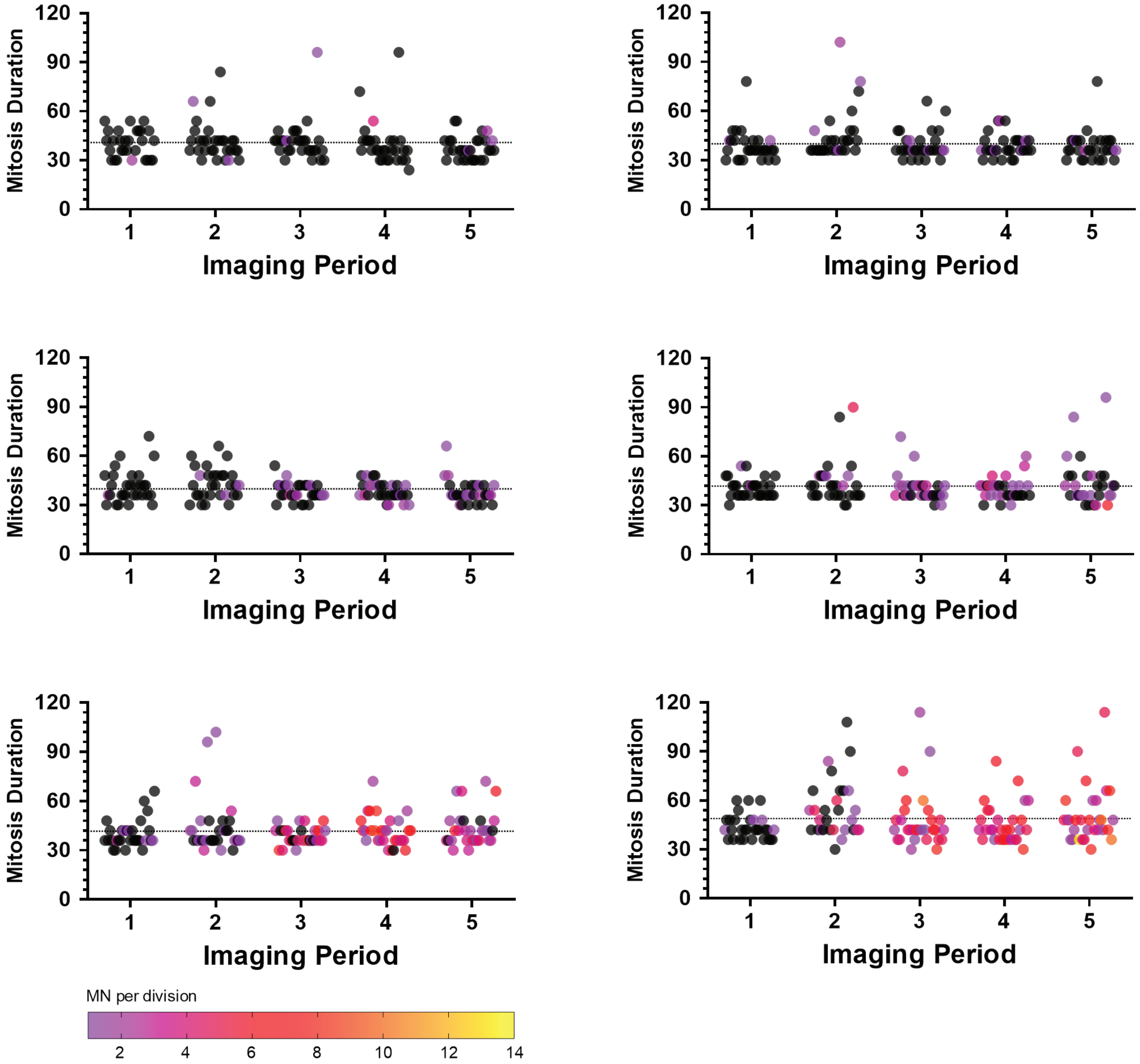
